# The Prmt5-Vasa module is essential for dichotomous spermatogenesis in *Bombyx mori*

**DOI:** 10.1101/2022.06.17.496537

**Authors:** Xu Yang, Dongbin Chen, Shirui Zheng, Meiyan Yi, Yongjian Liu, Yongping Huang

## Abstract

Spermatogenesis is a key process for the sexual reproduction species. In lepidopteran insects, dichotomous spermatogenesis is a notable feature, which produces eupyrene (nucleate) and apyrene (anucleate) spermatozoa. Both sperm morphs are essential for fertilization, as eupyrene sperm fertilizes the egg, while apyrene sperm is necessary for eupyrene sperm migration. In *Drosophila*, Prmt5 acts as a type II arginine methyltransferase to catalyze the symmetrical dimethylation of arginine residues (sDMA) in Vasa. However, Prmt5 is involved in regulating spermatogenesis, but Vasa is not. To date, the functional genetic study on dichotomous spermatogenesis in the lepidopteran model *Bombyx mori* has been limited, thus the underlying mechanism remains largely unknown. In this study, we report that both BmPrmt5 and BmVasa act as essential components in the regulation of dichotomous spermatogenesis in *Bombyx mori*. The loss-of-function mutants of *BmPrmt5* (*ΔBmPrmt5*) and *BmVasa* (*ΔBmVasa*) derived from CRISPR/Cas9-based gene editing show similar male and female-sterile phenotypes. Through immunofluorescence staining analysis, we found that the morphs of both *ΔBmPrmt5* and *ΔBmVasa* sperm show severe defects, indicating an essential role for both BmPrmt5 and BmVasa in the regulation of dichotomous spermatogenesis. RNA-seq analyses indicate that the defects in dichotomous spermatogenesis observed for *ΔBmPrmt5* and *ΔBmVasa* mutants could be attributed to the reduced expression of the spermatogenesis-related genes including *Sex-lethal* (*BmSxl*), implying that BmSxl may act downstream of Prmt5 and Vasa in regulating apyrene sperm development. These findings suggest that BmPrmt5 and BmVasa constitute an integral regulatory module essential for dichotomous spermatogenesis in *Bombyx mori*, in which BmPrmt5 may promote BmVasa activity through sDMA modification.

**Author Summary:** Dichotomous spermatogenesis is a notable feature in Lepidoptera insects, in which a single male produces nucleated and anucleated dimorphic sperm. Both sperm morphs are essential for fertilization, as the eupyrene sperm fertilizes the egg, while the apyrene sperm assists in transporting the eupyrene sperm to the female sperm storage organs. To date, the mechanism of dichotomous spermatogenesis is still unclear, as very limited components participating in this process have been reported. Here we defined the function of BmPrmt5 (Protein arginine methyltransferase 5) and its substrate BmVasa in dichotomous spermatogenesis in the lepidopteran insect *Bombyx mori*. Our genetic analyses revealed that both *BmPrmt5* and *BmVasa* are essential for dichotomous spermatogenesis. Moreover, RNA-seq analyses, in conjunction with the genetic and immunofluorescence staining analyses, suggest a potential regulatory pathway consisting of the BmPrmt5–BmVasa module in dichotomous spermatogenesis in *Bombyx mori*. This study expands the knowledge of reproduction biology in insects, and may provide potential gene targets for pest control.

## Introduction

Spermatogenesis is a vital process for the sexual reproduction species [1]. During spermatogenesis, the germ cells undergo mitosis and meiosis and complete morphogenetic changes [2,3]. Despite the central role of spermatozoa in reproduction, it exhibits exceptional diversities at both ultrastructure and molecular levels [4,5], thus becomes a valuable model for molecular genetics research. Up to date, the studies on the mechanism of spermatogenesis have largely been limited to the model species [6], which could not reflect the natural diversities from an evolutionary perspective.

Lepidopteran is an emerging and notable taxon for spermatogenesis research. Understanding the reproductive processes in Lepidoptera is essential because it includes both pest species and species of economic importance [7,8]. Besides, sperm dichotomy is an impressive phenomenon present throughout Lepidoptera [9], excluding only two species from the primitive Micropterigidae [10]; however, it is absent not only in Drosophila, but also in the sister order, Trichoptera [11], and the two closely related orders, Diptera and Siphonaptera [12]. Like most other Lepidopterans, the Lepidoptera model *Bombyx mori* displays dichotomous spermatogenesis characterized by the production of eupyrene (nucleate) and apyrene (anucleate) spermatozoa [13]. Both sperm morphs are essential for fertilization, as eupyrene sperm fertilizes the egg, while apyrene sperm assists in transporting eupyrene sperm to the female sperm storage organs [14,15]. To date, the studies on dichotomous spermatogenesis have primarily been limited to the aspects related to cytology and developmental timing by microscopic observations, and only few genes such as *poly(A)-specific ribonuclease-like domain-containing 1* (*BmPnldc1*), *Sex-lethal* (*BmSxl*), and *Maelstrom* (*BmMael*) have been experimentally linked to this process in *B. mori* [14–20]. Therefore, the molecular and genetic framework that controls this process remains largely unknown.

Post-translational modifications (PTMs) play critical roles in diverse cellular events, including DNA damage response, chromosome condensation, and cytoskeletal organization during germ cell differentiation [21,22]. It is known that *Protein arginine methyltransferase 5* (*Prmt5*) participates in PTM, which is a type II arginine methyltransferase that catalyzes the symmetrical dimethylation of arginine residues (sDMA) of its substrates [23,24]. Methylated arginines in particular sDMAs bind to Tudor domains of proteins (Tudors) and regulate protein-protein interaction and protein cellular localization [25–27]. Prmt5 plays important roles in varied developmental processes among species [24,28,29]. For instances, it is required for early embryonic development and stem cell differentiation in mouse[30–32]; In *Drosophila melanogaster*, mutation of *Dart5* homologous to *Prmt5* results in female grandchildless and male spermatogenesis defects, and disruption of the circadian rhythms in locomotor activity [33–36]; In zebrafish, loss function of *Prmt5* causes a reduction in germ cell number, leading to the failure of gonads to differentiate into normal testis or ovaries and eventual sterility [37]. Whether Prmt5 plays a role in regulating dichotomous spermatogenesis in *Bombyx mori* is not known.

A number of Prmt5 substrates such as Vasa, P-element-induced wimpy testis proteins (PIWIs) and Tudors have been identified through biochemical and genetic studies [26,38], which play an essential role in regulating germ cell specification in *Drosophila, Xenopus*, and mouse [21]. Of them, Vasa is a member of the DEAD-box family protein conserved in the phyla that functions as an ATP-dependent RNA helicase [39], which is specifically expressed in the germline in these species [40,41]. The Xenopus Vasa homolog is expressed in oocytes and embryos and is required for the formation of germ cells [42–44]. Mouse Vasa homolog MVH (also known as DDX4) expression is also restricted to the germ cell lineage, and its loss of function causes a deficiency in the proliferation and differentiation of spermatocytes [45]. In *Drosophila*, Vasa is required for the assembly and function of the pole plasm during oogenesis [46,47]. Like Prmt5, whether Vasa functions in spermatogenesis in insects other than *Drosophila* is also unknown.

In this study, we characterized the functions of BmPrmt5 and BmVasa in regulating dichotomous spermatogenesis in *B. mori*. We first generated the loss-of-function mutant of *BmPrmt5* (*ΔBmPrmt5*) through CRISPR/Cas9-based gene editing, found that it displays severe defects during dichotomous spermatogenesis, and male and female sterility. We show by immunofluorescence staining assay of protein cellular localization that BmVasa is expressed in sperm cysts throughout spermatogenesis. We then constructed the loss-of-function mutant of *BmVasa* (*ΔBmVasa*) through CRISPR/Cas9-based gene editing, and found that this mutant also shows severe defects in dichotomous spermatogenesis, and is male- and female-sterile, similar to *ΔBmPrmt5* mutant. To gain insights into the changes in global gene expression associated with spermatogenesis defects in *ΔBmPrmt5* and *ΔBmVasa* mutants, we performed RNA-seq analysis, and found that numerous cell-differentiation-related genes are significantly decreased. The combined results from the genetic, immunofluorescence staining, and RNA-seq analyses demonstrate that the defects in eupyrene and apyrene sperm observed for *ΔBmPrmt5* and *ΔBmVasa* mutants could be attributed to disorganized cellular structural proteins and cell polarity disorder. This study suggests that BmPrmt5 and BmVasa act as an integral regulatory module essential for dichotomous spermatogenesis in *B. mori*.

## Results

### Mutation of *BmPrmt5* leads to both female and male sterility in *Bombyx mori*

Previous study have shown that Prmt5 is predominantly expressed in the germline of *Drosophila* [33]. To determine the possible organ-specific expression pattern of *BmPrmt5* in *Bombyx mori* (*B. mori*), we performed qRT-PCR analysis with the samples prepared from different organs and found that *BmPrmt5* was predominantly expressed in the gonads at different stages (Fig 1A). To verify *BmPrmt5* function in the gonads, we first used a binary transgenic CRISPR/Cas9 system to obtain its loss-of-function mutant, *ΔBmPrmt5*. To this end, we designed two small guide RNAs (sgRNAs) targeting exons 1 and 2 of *BmPrmt5*, and *ΔBmPrmt5* mutants were generated through genetic crossing between the U6-sgRNA lines and the nos-Cas9 lines (Fig 1B). Mutations in the randomly selected representative F1 offsprings were detected by genomic PCR and DNA sequencing, which indicates that the ORF of *BmPrmt5* was shifted in both male and female *ΔBmPrmt5* individuals (Fig 1B). Further qRT-PCR analysis showed that *BmPrmt5* was hardly expressed in *ΔBmPrmt5* (Fig 1C). The *ΔBmPrmt5* adults were viable and grossly normal, and the *ΔBmPrmt5* females laid significantly fewer eggs than the wild-type (WT). (Fig 1D and E). We then performed fecundity testis, and found that the crossing between *ΔBmPrmt5* female and WT male or between *ΔBmPrmt5* male and WT female generated the hybrids showing both female and male sterility (Fig 1F). These results demonstrate that BmPrmt5 is essential for both female and male fertility.

**Fig 1.**
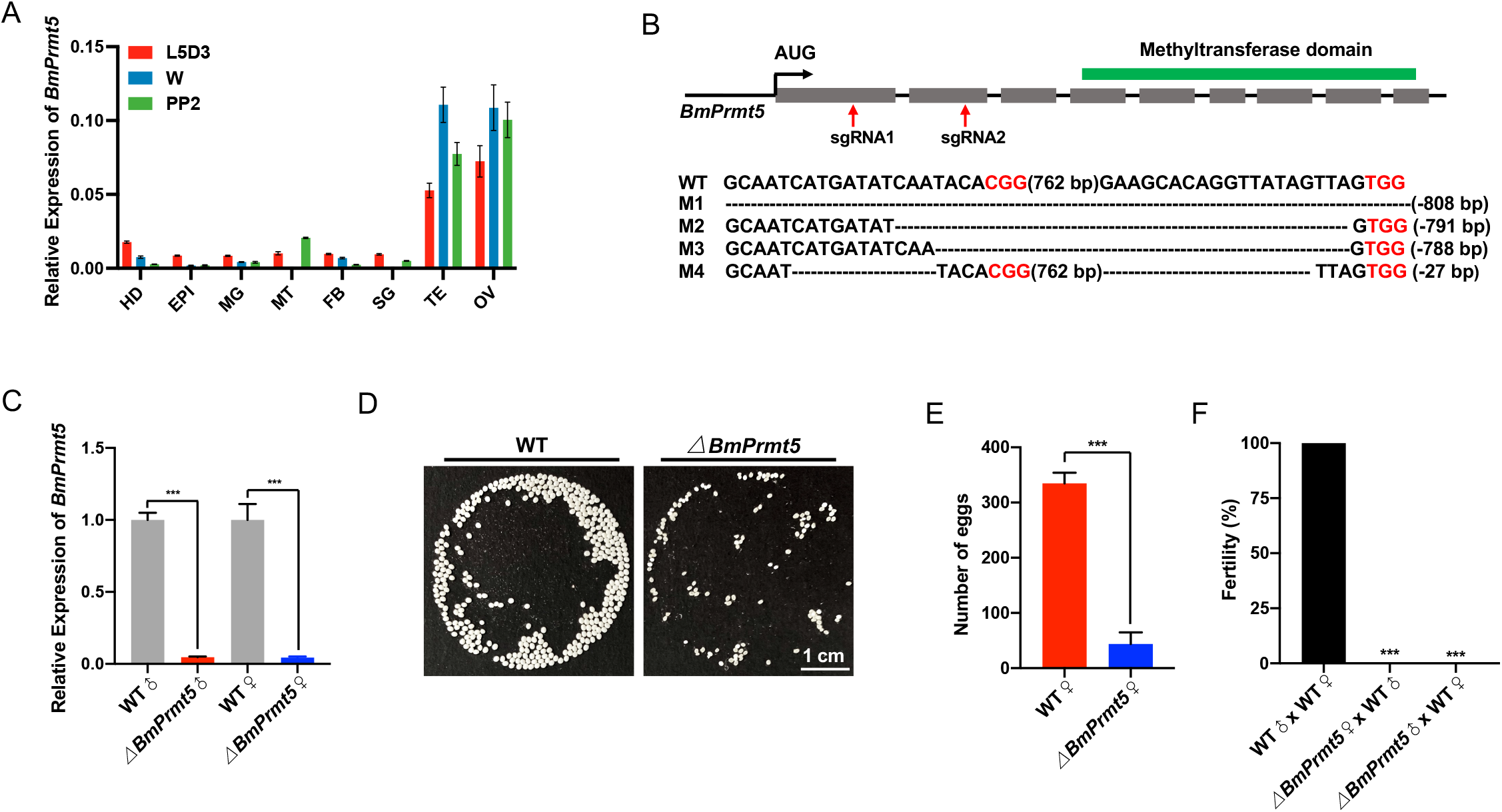
Mutation of *BmPrmt5* causes both male and female sterility in *Bombyx mori*. (A) qRT-PCR analysis of of *BmPrmt5* mRNA levels in eight tissues at three different stages. HD, head; EPI, epidermis; MG, midgut; MT, malpighian tubule; FB, fat body; SG, silk gland; TE, testis; OV, ovary; L5D3, third day of the fifth larval stage; W, wandering stage; PP2, the second day of the prepupal stage. *Ribosomal protein 49* (*Ro49*) was used as an internal reference. Data are mean ± SEM. (B) Genomic disruption of the *BmPrmt5* gene using CRISPR/Cas9 schematic of *BmPrmt5* gene structure and sgRNA targets. Gray filed boxes represent exons of the ORFs, and the function domain predicted by SMART is highlighted with a green box above. Red arrows indicate the target sites of sgRNA1 and sgRNA2. Genomic mutations of the *BmPrmt5* gene are shown with target sequences. The dashed lines indicate deleted sequences. M1-4, represents four mutation lines. (C) Transcript levels of *BmPrmt5* detected by qRT-PCR in testis and ovary of WT and *ΔBmPrmt5* at the wandering stage. Three individual biological replicates were performed, and the error bars are mean ± SEM. The asterisks (***) indicate significant differences relative to WT (*P* < 0.001, t-test). (D) Images of eggs laid by WT and *ΔBmPrmt5*. Scale bar, 1 cm. (E) The number of eggs laid by WT and *ΔBmPrmt5* (n=6, *P* < 0.001, t-test). (F) Fertility of males and females of the indicated genotypes. Fertility is indicated on the histogram (n= 15, *P* < 0.001, Fisher exact test)

### BmPrmt5 is essential for dichotomous spermatogenesis

The demonstration that *ΔBmPrmt5* shows male sterility indicates possible defects in the reproduction system (Fig 1F). To confirm this possibility, we first investigated whether the *ΔBmPrmt5* males would exhibit any apparent defects in the genitalia or the reproductive system. However, no obvious defects in the genitalia or the reproductive system were detected. It is known that the eupyrene and apyrene sperms of *B. mori* show different morphology and timing of differentiation during spermatogenesis [17–19]. Specifically, the eupyrene spermatogenesis begins on the first two days at the fifth instar larval stage, while the apyrene spermatogenesis starts during the wandering stage. We then explored whether BmPrmt5 would be required for spermatogenesis. To do this, we first performed fluorescence staining to examine the development of both eupyrene and apyrene sperm bundles on the seventh day at the pupal stage in WT and *ΔBmPrmt5*. The results showed that the needle-shaped sperm nuclei were assembled regularly at the head of the eupryrene sperm bundles in the WT, whereas *ΔBmPrmt5* eupryrene sperm nuclei were abnormally organized and exhibited squeezed eupyrene sperm bundles (Fig 2A). As for the apyrene sperm, WT cysts of the elongated apyrene spermatids had no polarity, with small round micronuclei being distributed in the middle region (Fig 2B). By contrast, almost all the *ΔBmPrmt5* apyrene sperm bundles exhibited defects in sperm nucleus shape and localization, and only few normal apyrene sperms existed in the testis at the late pupae stage in *ΔBmPrmt5*.

**Fig 2.**
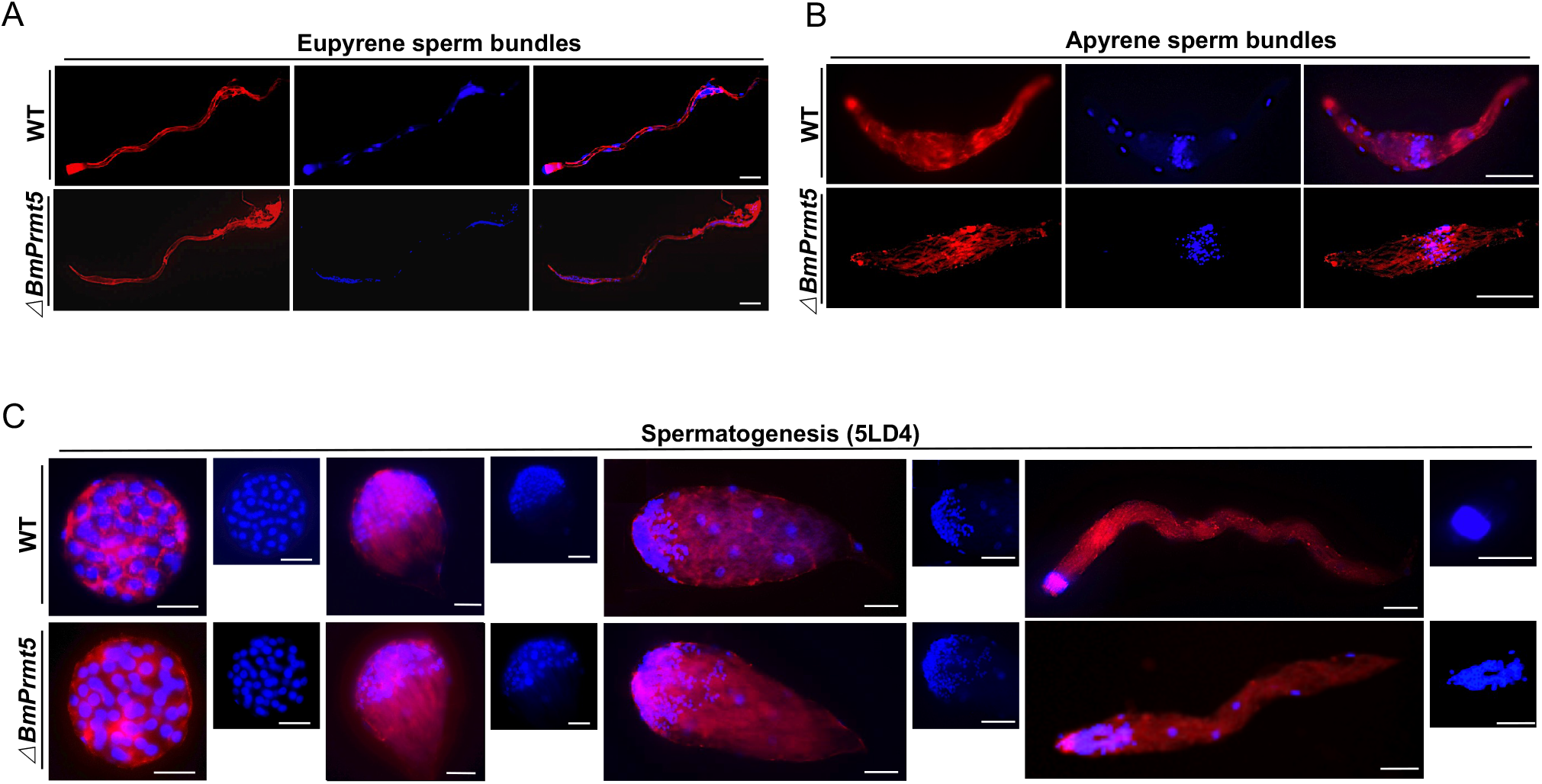
Loss-of-function of *BmPrmt5* leads to defects in spermatogenesis. (A and B) Representative immunofluorescence images of eupyrene sperm bundles (A) and apyrene sperm bundles (B) from WT and *ΔBmPrmt5* on the seventh day of the pupal stage. Scale bars, 50 μm. (C) Representative immunofluorescence images of eupryrene sperm bundles at different developmental stages at day four of the fifth larval instar of WT and *ΔBmPrmt5*. Scale bars, 50 μm. Blue, Hoechst; red, F-actin.

Next, we investigated eupyrene spermatogenesis in the testes on the fourth day of the fifth larval instar in WT and *ΔBmPrmt5* in details. The results showed that, at the early elongating stage, *ΔBmPrmt5* developed similar eupyrene sperm bundles to WT with round nuclei being localized at the anterior part of the bundles (Fig 2C). At the late elongating stages, however, the eupyrene sperm nuclei were severely dislocated in *ΔBmPrmt5*, but not in WT (Fig 2C). Taken together, these results demonstrate that BmPrmt5 plays a vital role in the regulation of dichotomous spermatogenesis.

### BmVasa is expressed in spermatocytes and sperm bundles throughout spermatogenesis in *Bombyx mori*

It is reported that Prmt5 specifically catalyzes symmetric dimethylation of the arginine residues (sDMA) in its substrates including Vasa to regulate their activity in gonad tissues among the species across phyla [35–37,48]. We first analyzed the putative sDMA motifs of BmVasa by aligning Vasa homologs in lepidopterans and the well-identified species including *Caenorhabdits elegans, Danio rerio, Xenopus laevis, Mus musculus*, and *Drosophila melanogaster* (S1 Fig). The results demonstrate that there exists a putative conserved sDMA motif in the N terminal of BmVasa (Fig 3A). To explore whether the Prmt5 and Vasa might act as a module to regulate spermatogenesis in the Insecta, we then verified the expression patterns of *BmPrmt5* and *BmVasa* in testis at the different stages during the spermatogenesis in *B. mori* by qRT-PCR. The results showed that the expression of both *BmPrmt5* and *BmVasa* started to increase on the fifth day at the fifth larval stage and peaked at the wandering stage, and then gradually decreased till the third day at the prepupa stage (Fig 3A and B). We then used the antibody against Vasa to detect the cellular localization of BmVasa in spermatocytes by immunohistochemistry and found that it was localized in the cytoplasm (S4 Fig). We also observed that BmVasa was expressed in sperm cysts throughout different development stages (Fig 3D). These results indicate that BmVasa is expressed in the sperm bundles during spermatogenesis.

**Fig 3.**
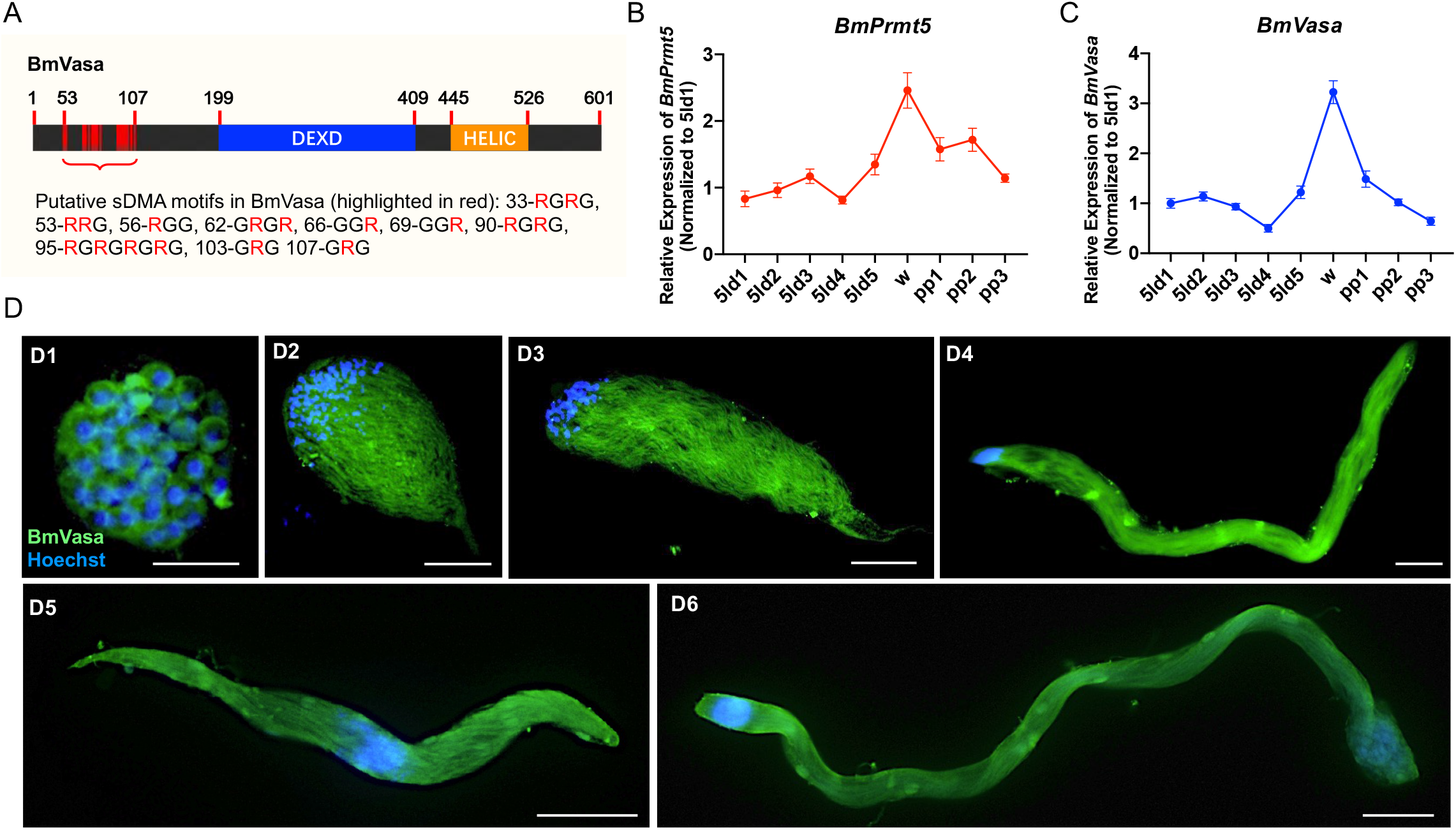
*BmVasa* is expressed in spermatocytes and sperm bundles throughout spermatogenesis in *Bombyx mori*. (A) Schematic of BmVasa and its putative symmetric dimethylated arginines residues (sDMA). Numbers refer to amino acid positions. DEXD, DEAD-like helicases superfamily domain; HELIC, helicase superfamily C-terminal domain. (B and C) qRT-PCR analyses of the transcript levels of *BmPrmt5* (B) and *BmVasa* (C) in testis from day one of the fifth instar to the third day of pupa stage. (D) Representative immunofluorescence images of spermatocytes and sperm bundles. Spermatocytes (D1) and elongating eupyrene sperm bundles (D2-D4) on the fourth day of the fifth instar larval stage; Apyrene sperm bundles (D5) and eupyrene sperm bundles (D6) on the seventh day of pupa stage. Blue, Hoechst; Green, BmVasa. Scale bars, 50 μm.

### Mutation of *BmVasa* also leads to both female and male sterility

To further determine BmVasa participation in spermatogenesis, we first detected the expression pattern of *BmVasa* by qRT-PCR analysis with the samples prepared from different organs and found that it was predominantly expressed in the gonads at different stages (Fig 4A). We then created the loss-of-function mutants of *BmVasa* (*ΔBmVasa*) using CRISPR/Cas9 system. Two sgRNAs targeting exons 3 and 7 of *BmVasa* were designed, and *ΔBmVasa* mutants were generated by sexual crossing between the U6-sgRNA lines and the nos-Cas9 lines (Fig 4B). Mutations in randomly selected representative F1 offspring were detected by genomic PCR and sequencing using gene-specific primers, confirming that the mutations leading to the ORF shift of *BmVasa* were produced in both male and female individuals (Fig 4B). Further qRT-PCR and Western blotting analyses showed that either the *BmVasa* transcript or the BmVasa protein was hardly detectable in *ΔBmVasa* (Fig 4C and S3 Fig). The *ΔBmVasa* adults were viable and grossly normal, and the *ΔBmVasa* females laid significantly fewer eggs than WT (Fig 4D and E). We then performed fecundity testis and found that the crossing between *ΔBmVasa* female and WT male or between *ΔBmVasa* male and WT female generated the hybrids showing both female and male sterility (Fig 4F). These results demonstrate that, like BmPrmt5, BmVasa is also essential for both female and male fertility.

**Fig 4.**
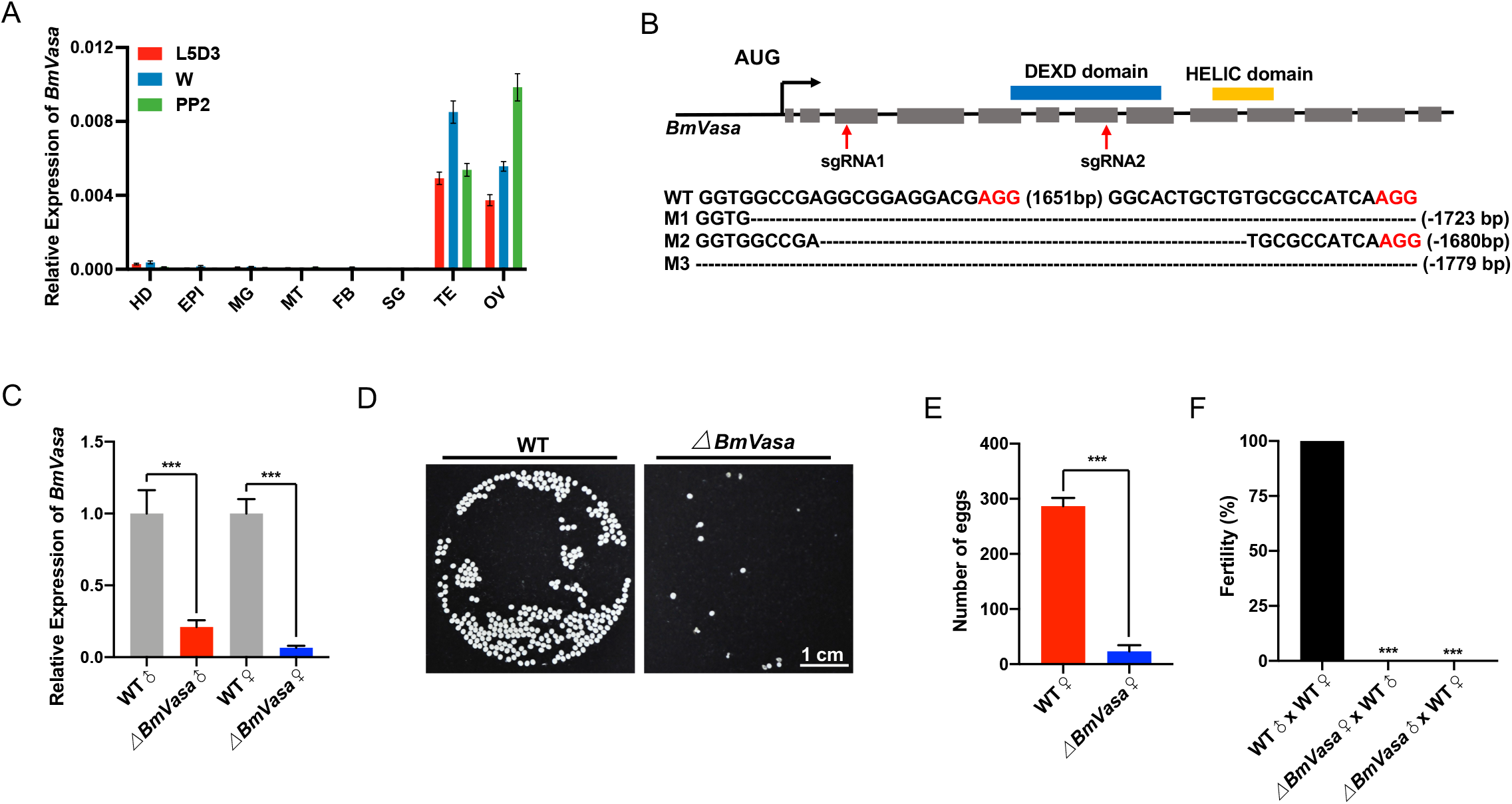
Loss-of-function of *BmVasa* causes both male and female sterility in *Bombyx mori*. (A) qRT-PCR analysis of *BmVasa* mRNA levels in eight tissues at three different stages. HD, head; EPI, epidermis; MG, midgut; MT, malpighian tubule; FB, fat body; SG, silk gland; TE, testis; OV, ovary; L5D3, third day of the fifth larval stage; W, wandering stage; PP2, the second day of the prepupal stage. *Ribosomal protein 49 (Rp49*) was used as an internal reference. Data are mean ± SEM. (B) Genomic disruption of the *BmVasa* gene using CRISPR/Cas9. Schematic of BmVasa gene structure and sgRNA targets. Gray filed boxes represent exons of the ORFs, and the function domains (DEXD and HELIC) predicted by SMART are highlighted with blue and yellow boxes above. Red arrows indicate the target sites of sgRNA1 and sgRNA2. Genomic mutations of *BmVasa* gene are shown with target sequences. M1-3, represents four mutation lines. The dashed lines indicate deleted sequences. (C) qRT-PCR analysis of the transcript levels of *BmVasa* in testis and ovary of WT and *ΔBmVasa* at the wandering stage. Three individual biological replicates were performed, and the error bars are mean ± SEM. The asterisks (***) indicate significant differences relative to WT (*P* < 0.001, t-test). (D) Images of eggs laid by WT and *ΔBmVasa* mutant. (E) The number of eggs laid by WT and *ΔBmVasa* (n=6, *P* < 0.001, t-test). (F) Fertility of males and females of the indicated genotypes. Fertility is indicated on the histogram (n= 15, *P* < 0.001, Fisher exact test)

### BmVasa is also required for dichotomous spermatogenesis

The demonstration that *ΔBmVasa* shows male fertility indicates defects in the reproduction system (Fig 4F). To further explore how BmVasa might affect spermatogenesis, we first detected the genitalia or the reproductive system in the *ΔBmVasa* males, and then examined the development of both eupyrene and apyrene sperm bundles on day seventh at the pupal stage in the *ΔBmVasa* males by fluorescence staining. Both *ΔBmVasa* eupyrene and apyrene sperm bundles showed severe abnormalities. Specifically, like *ΔBmPrmt5, ΔBmVasa* sperm nuclei exhibited squeezed eupyrene sperm bundles with abnormally organized sperm nuclei (Fig 5A). Moreover, like *ΔBmPrmt5*, all the *ΔBmVasa* apyrene sperm bundles had severe defects in sperm nucleus shape and location, and some of the defective sperm bundles displayed intermediate morphology between normal apyrene sperm and early elongating eupyrene sperm bundles (Fig 5B).

**Fig 5.**
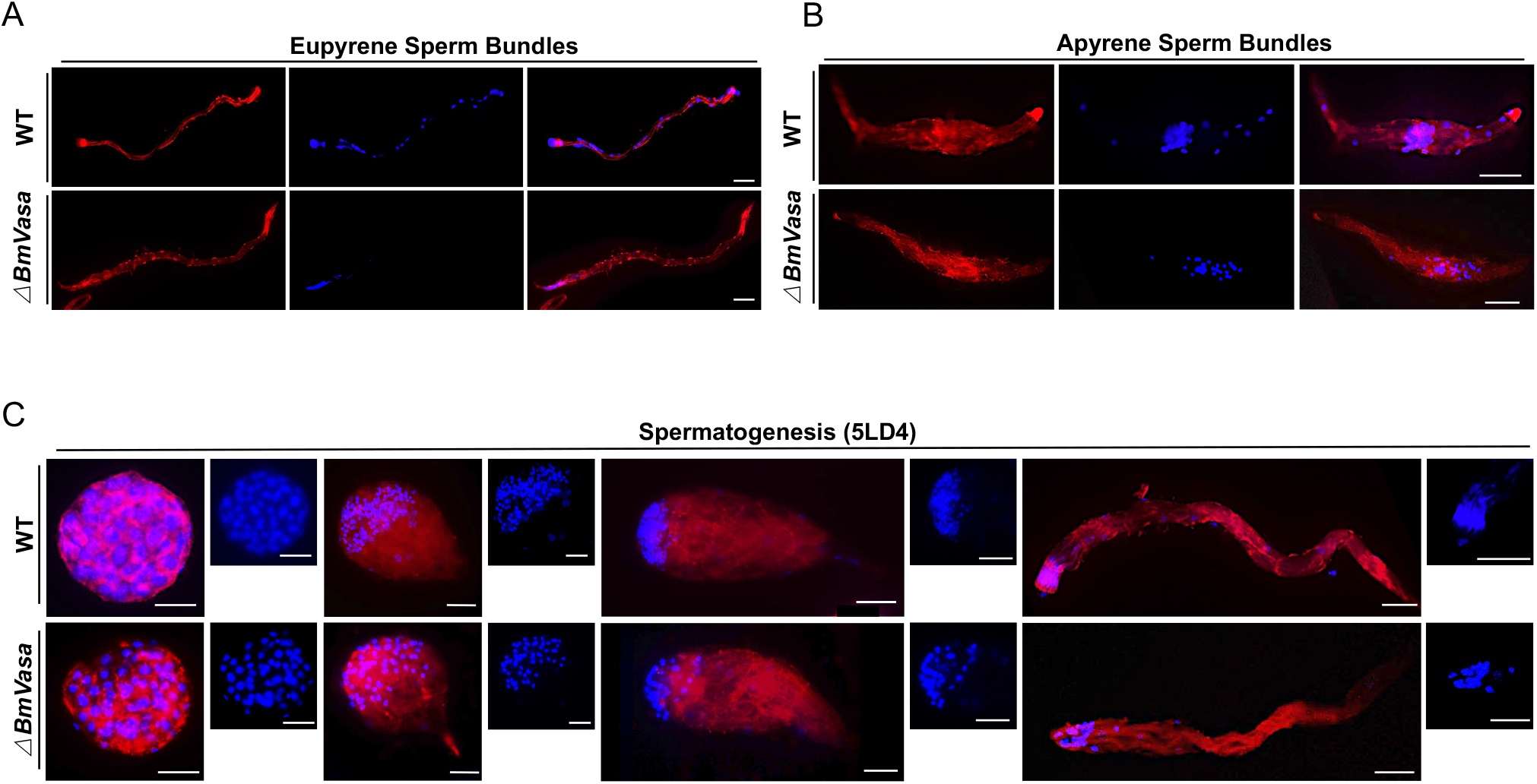
*BmVasa* mutation results in spermatogenesis defects. (A and B) Representative immunofluorescence images of eupyrene sperm bundles (A) and apyrene sperm bundles (B) from WT and *ΔBmVasa* on the seventh day of the pupal stage. Scale bars, 50 μm. (C) Representative immunofluorescence images of eupryrene sperm bundles in different developmental stages at day four of the fifth larval instar of WT and *ΔBmVasa*. Scale bars, 50 μm. Blue, Hoechst; red, F-actin.

Next, we investigated eupyrene spermatogenesis in the testis on day fourth of the fifth larval instar in detail using *ΔBmVasa*. The results showed that, like *ΔBmPrmt5*, at the early elongating stage, *ΔBmVasa* eupyrene sperm bundles were similar to WT, with the round nuclei being localized in the anterior part of the bundles. In the late elongating stages, however, like *ΔBmPrmt5*, the *ΔBmVasa* nuclei were severely dislocated (Fig 5C). Taken together, these results demonstrate that, like BmPrmt5, BmVasa is also required for dichotomous spermatogenesis.

### BmPrmt5 does not affect the transcript and protein level of BmVasa

To determine whether BmPrmt5 might influence the expression of *BmVasa*, we first performed qRT-PCR analysis using *ΔBmPrmt5* testis and sperm to exclude the possibility that BmPrmt5 would regulate the transcription of *BmVasa*. As anticipated, no significant changes in *BmVasa* expression were observed in WT and *ΔBmPrmt5* (S4A and B Fig). We then determined the levels of BmVasa protein in WT and *ΔBmPrmt5* by Western bolting using an anti-Vasa antibody, respectively. The results showed that BmVasa was expressed at a similar level in these two backgrounds (S4C Fig). These results indicate that BmPrmt5 does not affect BmVasa stability.

### BmPrmt5 and BmVasa co-regulate a large number of genes in the same direction

To gain insights into the changes in global gene expression associated with the defects in male fertility observed in *ΔBmPrmt5* and *ΔBmVasa* mutants, we first performed RNA-sequencing (RNA-seq) assays using spermatocytes and sperm bundles isolated from testis of WT, *ΔBmPrmt5*, and *ΔBmVasa* mutants, respectively. Up to 404 and 522 sperm-specific differentially expressed genes (DEGs) regulated by BmPrmt5 and BmVasa were identified, respectively (S5 Fig). Among them, 303 DEGs were co-regulated by BmPrmt5 and BmVasa, of which 238 (78%) and 60 (19%) were up-regulated and down-regulated by BmPrmt5 and BmVasa, respectively (Fig 6A). Next, we performed gene ontology (GO) enrichment analysis, and found that the genes regulated by BmPrmt5 and BmVasa were preferentially associated with gap junction, components of the extracellular region, and germ-line cyst formation (Fig 6B). After the unannotated and transposon-related DEGs co-regulated by BmPrmt5 and BmVasa were filtered, a heat map was generated by hierarchical clustering analyses, which reveals that 35 of these genes were regulated by BmPrmt5 and BmVasa in the same direction in both testis and sperm (Fig 6C). Among the genes significantly down-regulated by BmPrmt5 and BmVasa, some are related to spermatogenesis, which include those encoding Sex-lethal (BmSxl), Tubulin Polymerization Promoting Protein (TPPP) family protein-like (TPPP-like), Tctex1 domain-containing protein 1-like (Tctex-like), Actin-chr1, and β-Tubulin. BmSxl is an RNA binding protein, whose mutation leads to severe defects in apyrene sperm development in *B. mori* [14,15]. The TPPP-like protein, TPPP2, is shown to be crucial for sperm morphogenesis in mice [49]. In *Drosophila*, the ortholog of the Tctex-1 acts as the dynein light chain, and its null mutant displays a diffused nuclear sperm cyst [50,51]. We then verified the expression of *BmSxl, TPPP-like, Tctex-like, Actin-chr1*, and *ß-Tubulin* in WT, *ΔBmPrmt5* and *ΔBmVasa* mutants through qRT-PCR, respectively. The results showed that all these genes expression was severely repressed in *ΔBmPrmt5* and *ΔBmVasa* mutants (Fig 6D), confirming that both BmPrmt5 and BmVasa are involved in promoting the expression of these genes.

**Fig 6.**
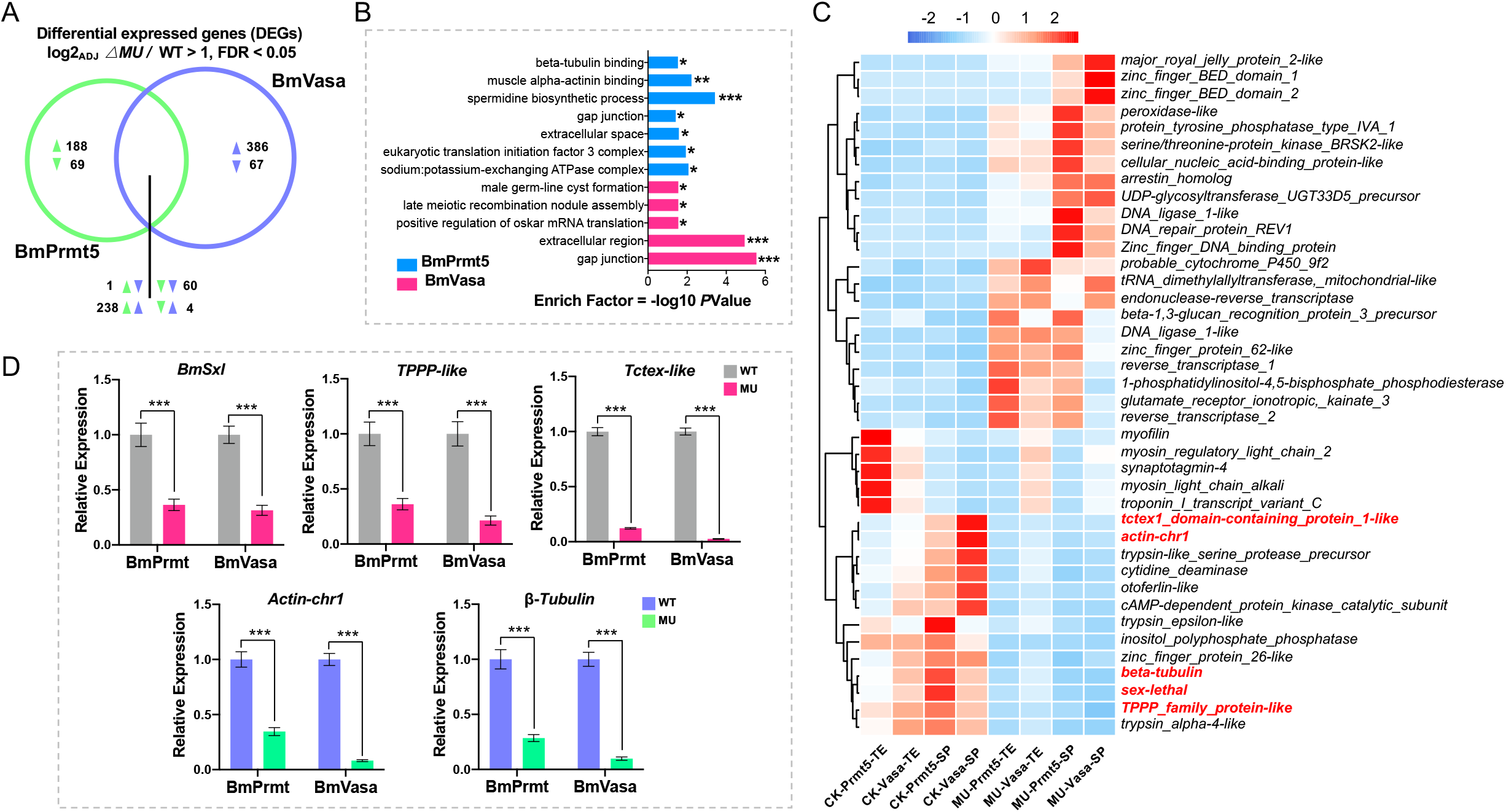
BmPrmt5 and BmVasa co-regulate many spermatogenesis-related genes in the same direction. (A) Venn diagram analysis showing sperm-specific DEGs commonly and specifically regulated by BmPrmt5 (left, green) and BmVasa (right, purple). DEGs, differentially expressed genes. (B) Gene ontology enrichment analysis showing the biological processes most associated with the sperm-specific DEGs regulated by BmPrmt5 and BmVasa. Enrichment analysis was conducted using the functional annotation tool DAVID. Statistical significance was determined at a p-value ≤ 0.05, *; p-value ≤ 0.01, **; p-value ≤ 0.001, ***; (C) Hierarchical clustering of selected spermatogenesis-related DEGs in different samples. CK-Prmt, control of BmPrmt5 mutant; CK-Vasa, control of BmVasa mutant; MU-Prmt, BmPrmt5 mutant; MU-Vasa, BmVasa mutant; TE, testis; SP, sperm. (D) qRT-PCR analysis of the selected spermatogenesis-related DEGs regulated by BmPrmt5 and BmVasa. Asterisks (***) indicates *P* < 0.001 determined by a two-tailed Student’s t-test. Data are mean ± SEM.

As BmSxl plays a critical role in regulating apyrene sperm development in *Bombyx mori* [14,15], we asked whether BmSxl might regulate the genes co-regulated by BmPrmt5 and BmVasa. To this end, we compared the 303 DEGs co-regulated by BmPrmt5 and BmVasa with the 528 BmSxl-regulated DEGs obtained previously [15], and found that 26 DEGs were co-regulated by BmPrmt5, BmVasa, and BmSxl (Fig 7A). A heatmap generated by hierarchical clustering analyses revealed that 9 DEGs including genes encoding TPPP family-like proteins were co-regulated by BmPrmt5, BmVasa and BmSxl in the same direction (Fig 7A). These results indicate that BmSxl may function in the same pathway of BmPrmt5 and BmVasa to regulate apyrene spermatogenesis (Fig 7B). Therefore, the defects in dichotomous spermatogenesis observed in *ΔBmPrmt5* and *ΔBmVasa* may be attributed to the reduced expression of spermatogenesis-related genes required for sperm morphogenesis.

**Fig 7.**
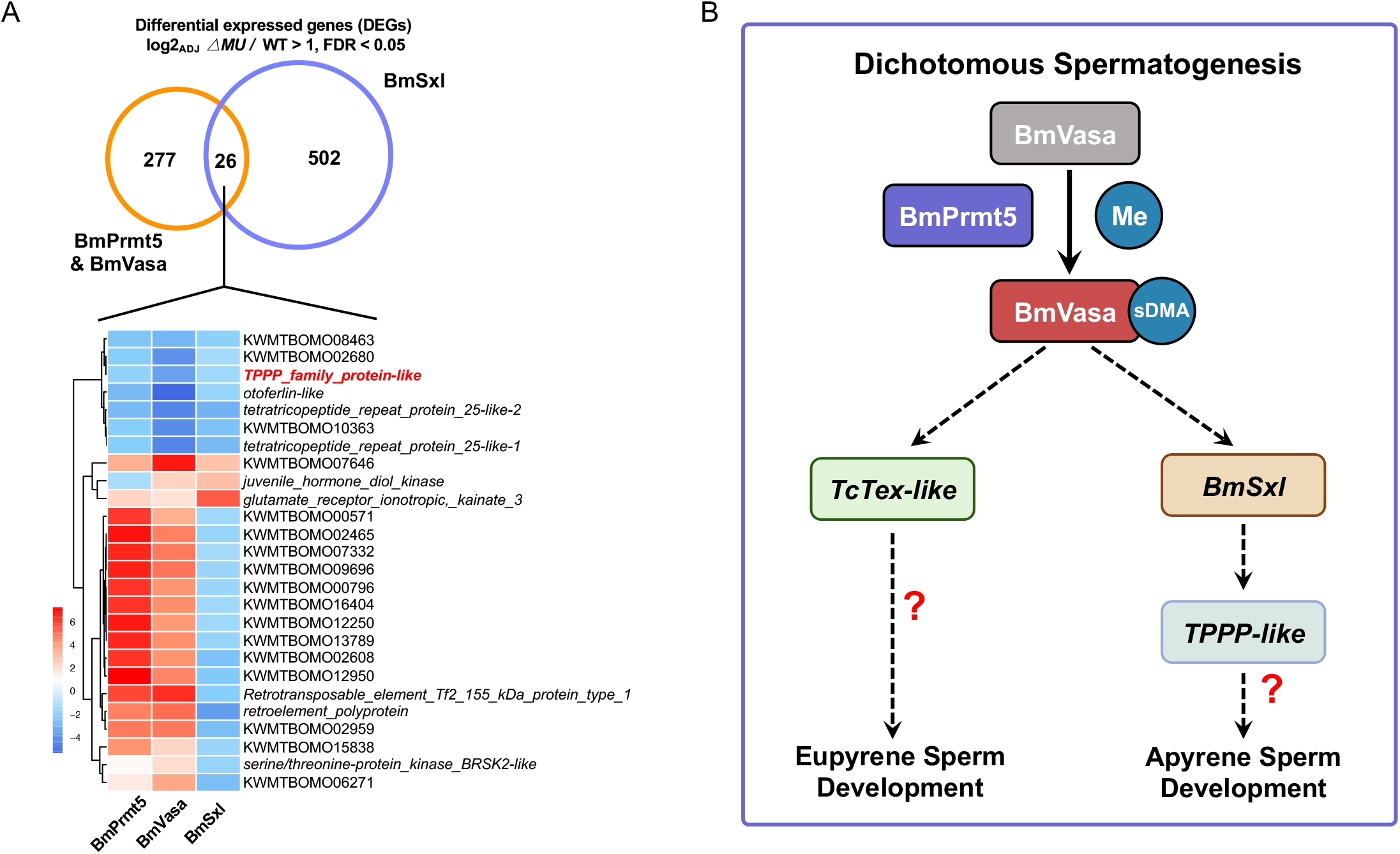
BmPrmt5, BmVasa, and BmSxl may act in the same pathway to regulate dichotomous spermatogenesis in *B. mori*. **(A)** Conjoint RNA-seq analysis with sperm cysts transcriptome data showing the DEGs regulated by BmPrmt5/BmVasa and BmSxl. Venn diagram analysis (upper panel) shows the DEGs commonly and specifically regulated by BmSxl (right, purple) and BmPrmt5/BmVasa (left, orange). Hierarchical clustering analysis (lower panel) shows the common DEGs regulated by BmPrmt5, BmVasa, and BmSxl. The fold change of each gene in different samples is shown in the heatmap (blue, down-regulated; red, up-regulated). **(B)** A model illustrating how the BmPrmt5-BmVasa module may regulate dichotomous spermatogenesis in *B. mori*. BmPrmt5 may catalyze the symmetrical dimethylation of arginine residues (sDMA) in BmVasa and activate BmVasa. As a result, BmVasa is somehow able to enhance the expression of spermatogenesis-related genes such as *BmSxl, BmTPPP-like* and *BmTctex-like*, and eventually promote eupryrene and apyrene sperm development in *B. mori*.

## Discussion

It is known that in *Drosophila*, both Prmt5 and Vasa is essential for oogenesis [35,36]. However, whether Prmt5 and Vasa functions in spermatogenesis in insects other than *Drosophila* is still unknown. In our current study, we identified BmPrmt5 and BmVasa as new important components involved in the regulation of dichotomous spermatogenesis in *B. mori*. We generated the loss-of-function mutants of *BmPrmt5* and *BmVasa* through CRISPR/Cas9-based gene editing, respectively (Fig 1B and 5B). Phenotype analyses of these mutants demonstrate that both BmPrmt5 and BmVasa are required for both female and male fertility, and normal morphs of sperm bundles (Fig 1E and 4E). The combined results from protein immunohistochemistry and fluorescence staining assays demonstrate that BmPrmt5 and BmVasa play essentially an equal role in regulating dichotomous spermatogenesis in *B. mori*.

How may BmPrmt5 and BmVasa be equally required for dichotomous spermatogenesis *B. mori?* It has been established that Prmt5 catalyzes the modification of sDMA in PIWIs and Tudors in gonads among the species across phyla [25,36]. In *Drosophila*, mutation of Dart5 (homolog of Prmt5) leads to a failure in the sDMA modification of PIWIs and Tudors, which compromises their ability to interact with other proteins and to be properly localized in the cells, eventually resulting in severe defects in oogenesis [35,36]. However, our previous studies have shown that mutation of either PIWIs or Tudors basically barely affects male fertility in *B, mori* [52,53]. In this study, as mutation of BmPrmt5 does not affect the protein stability of BmVasa (S4C Fig), BmPrmt5-mediated sDMA modification of BmVasa may not regulate BmVasa protein stability but may activate BmVasa. In other words, only sDMA-modifed BmVasa may be active and able to regulate dichotomous spermatogenesis in *B. mori*. We therefore propose that BmPrmt5 and BmVasa act as an integral module in regulating dichotomous spermatogenesis in *B. mori* (Fig 7B).

Previous studies have demonstrated that morphology defects in the bundles of either eupyrene or apyrene sperm compromise their physiological function [14,16,54]. In our current study, we also demonstrate that both morphs of sperms display severe defects in *ΔBmPrmt5* and *ΔBmVasa* mutants. To explore how BmPrmt5 and BmVasa might work, we performed RNA-seq analysis and our results indicated that the eupyrene sperm sterility observed in *ΔBmPrmt5* and *ΔBmVasa* could be attributed to disorganized structural protein genes which are involved in spermatogenesis (Fig 6), as their phenotype is similar to *ΔLIS-1* and *ΔTctex-1* in *Drosophila* [50,51]. It is worth mentioning that the phenotype of eupyrene sperm in both *ΔBmPrmt5 and ΔBmVasa* obtained in this study also mimics that of *ΔBmPnldc1* (Fig 2A and 5A)[15], although *BmPnldc1* was somehow not identified in our RNAseq analysis. On the other hand, the defects in apyrene sperm characterized by abnormal polarized nuclei in *ΔBmPrmt5* and *ΔBmVasa* phenocopy those of *ΔBmSxl* (Fig 2B and 5B)[14,15]. Based on these previous reports and the results obtained from genetic, cellular, biochemical, and gene expression studies (this work), we propose that the BmPrmt5-BmVasa module may achieve its role in dichotomous spermatogenesis, at least in part, through BmSxl, TPPP-like, and TcTex-like (Fig 7B).

In this study, we have also revealed a nonconserved function of Vasa between Diptera and Lepidoptera. In *Drosophila*, Vasa is required for oogenesis and embryo development, but not for male fertility [46,47]. However, our results demonstrate that BmVasa is essential for both female and male fertility in *B. mori* (Fig 4F). Based on the limited knowledge in Tribolium, aphids, and bees [55–57], we are not sure whether the requirement of Vasa for male fertility during spermatogenesis is an exception in *B. mori* across Insecta. In *Drosophila*, a proposed mechanism by which Vasa controls germ cell formation is that it regulates the localization and translation of the proteins encoded by the pole plasm-related genes via its direct interactions with their RNAs and proteins [57–59]. As *ΔBmVasa* apyrene sperms display dislocalized polarized nuclei, some of which show an intermediate phenotype between eupyrene and apyrene sperm bundles (Fig 4B and C), BmVasa may be required for the pole plasm of apyrene sperm cyst development.

How may BmVasa regulate *BmSxl, TPPP-like*, and *TcTex-like* to regulate eupyrene sperm and apyrene development? We hypothesize that there may be transcription factors (TFs) that act independently to regulate the expression of *BmSxl* and *BmTctex-like*. Upon sDMA modification by BmPrmt5, BmVasa may get activated and be able to bind to the mRNAs of the genes encoding these TFs and manipulate their translation. As a result, they accumulate and regulate the expression of genes promoting eupyrene and apyrene sperm development such as *BmSxl* and *BmTctex-like*, and eventually promote dichotomous spermatogenesis in *B. mori*. It will be worth making efforts to look for these postulated TFs and investigate how they may be regulated by BmVasa in future studies.

## Materials and methods

### Silkworm strains

The multivoltine, non-diapausing silkworm strain, Nistari, was used in this study. Larvae were reared on fresh mulberry leaves under standard conditions at 25□.

### RNA isolation, cDNA synthesis, and quantitative real-time PCR (qRT-PCR)

Total RNA was extracted from three individual mutants or WT animals using the TRIzol reagent according to the manufacturer’s instructions (YEASEN). An aliquot of 1 μg of the total RNA was used to synthesize cDNA using PrimeScript™ RT reagent Kit with gDNA eraser (Takara). qRT-PCR analysis was performed on a StepOnePlus Real-Time PCR system (Applied Biosystems) with an SYBR green Real-Time PCR master mix (Toyobo). The *B. mori ribosomal protein 49 gene* (*Bmrp49*) was used as an internal control to standardize the variation of different templates. The amplification program was as follows: the samples were incubated at 95□ for 5 min, followed by 40 cycles of 95□ for 15 s and 60□ for 1 min. Sequences of the qRT-PCR primers are listed in the supporting information Table 1.

### Silkworm germline transformation and CRISPR/Cas9-mediated construction of *ΔBmPrmt5* and *ΔBmVasa*

A binary transgenic CRISPR/Cas9 system was used to construct *ΔBmPrmt5* and *ΔBmVasa*. The nos-Cas9 transgenic silkworm lines (nos-Cas9/IE1-EGFP) expressed the Cas9 nuclease under the control of the *B. mori* nanos promoter [60]. The plasmids for expression of two sgRNA targeting each gene (U6-sgRNA/IE1-DsRed) respectively were under the control of the U6 promoter were constructed to generate *ΔBmPrmt5* and *ΔBmVasa*. Primers for plasmid construction and sgRNA targeting sequences are listed in Table 1.

**Table 1.**
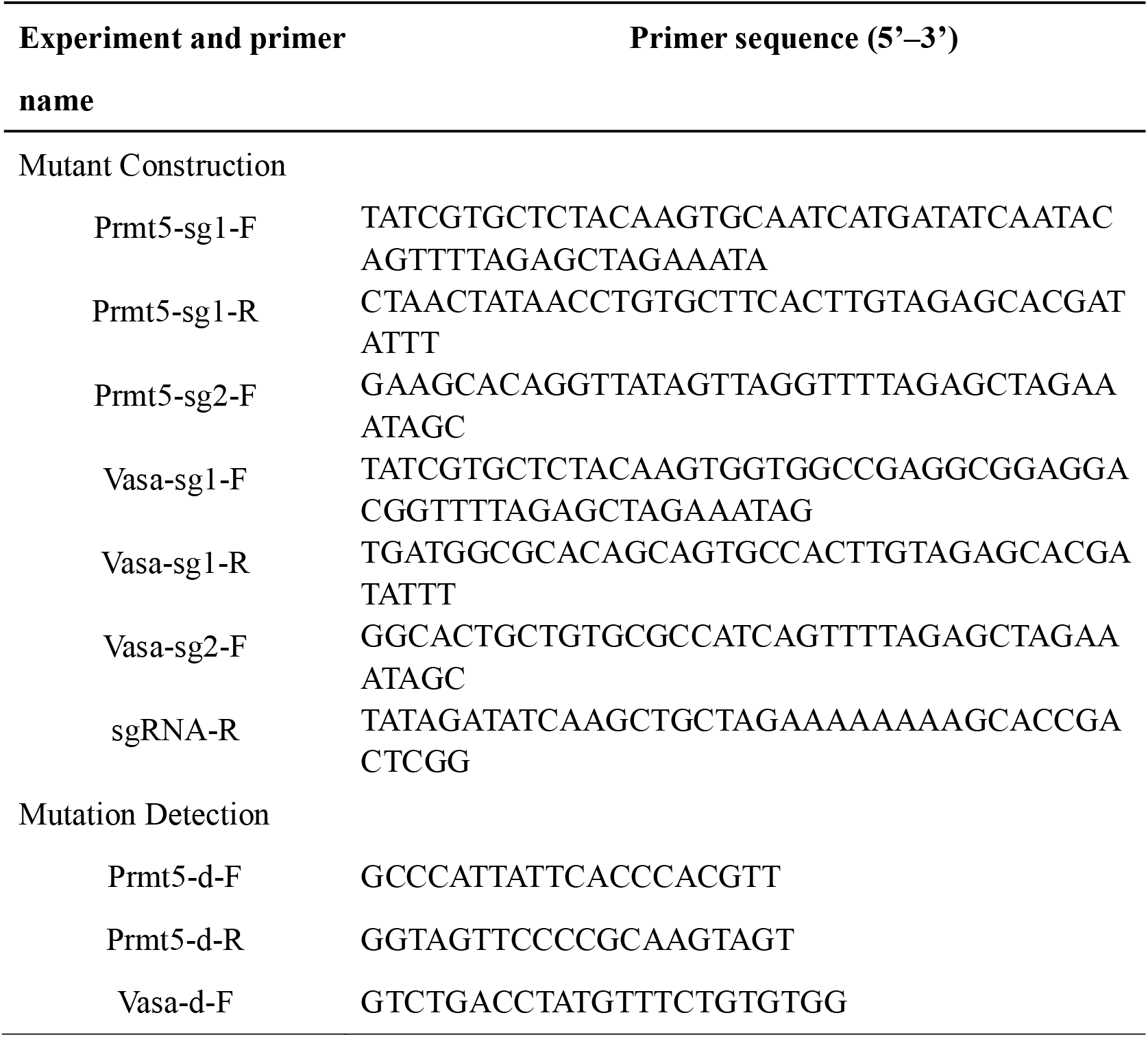
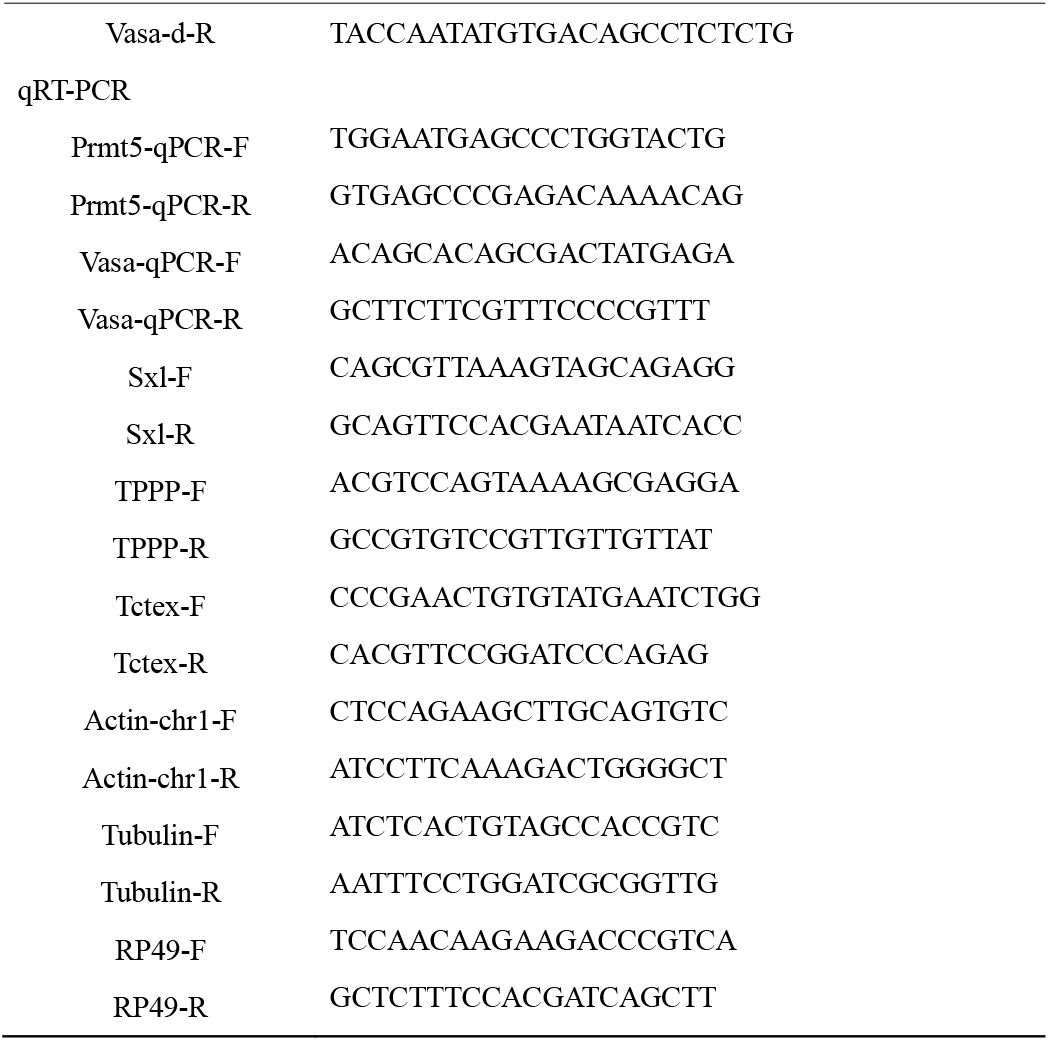
Primers used in this study

For silkworm germline transformation, preblastodermal embryos were prepared and microinjected with transgenic plasmids (400 ng/μl) together with helper plasmids (IFP2, 200 ng/μl) and incubated in a humidified chamber at 25□ for 10-12 days until hatching [61,62]. Putative transgenic generation 0 (G0) moths were sib-mated or backcrossed with wild-type moths, and G1 progeny were screened during early larval stages according to the GFP fluorescent marker using fluorescence microscopy (Nikon AZ100, Japan).

The nos-Cas9 lines and the U6-sgRNA lines were crossed to generate *ΔBmPrmt5* and *ΔBmVasa* with both EGFP and DsRed fluorescence markers for the subsequent experiments. Genomic DNA of the mutated animals was extracted by standard SDS lysis-phenol treatment, incubated with proteinase K, and purified for mutagenesis analysis via PCR amplification with specific primers (Supporting information Table 1).

### Phylogenetic and amino acid alignment analysis

Phylogenetic analysis and amino acid alignment analysis were performed by using MEGA 7 [63]. For phylogenetic analysis, evolutionary history was inferred using the neighbor-joining method [64]. The percentages of replicate trees in which the associated taxa clustered together in the bootstrap test (1000 replicates) were determined as previously described [65]. The evolutionary distances were computed using the Poisson correction method and are in units of the number of amino acid substitutions per site. Protein sequence of Vasa were download from NCBI (National Center for Biotechnology Information), the accession numbers are as follows: XP_037961717.1, *Plutella xylostella*; CAA72735.1, *Danio rerio*; EDL18409.1, *Mus musculus*; NP_001034520.2, *Tribolium castaneum*; NP_001037347.1, *Bombyx mori*; NP_001037347.1, *Manduca sexta*; NP_001081728.1, *Xenopus laevis*; NP_491113.1, *Caenorhabditis elegans*; NP_723899.1, *Drosophila melanogaster*; XP_001603956.3, *Nasonia vitripennis*; XP_013187571.1, *Amyelois transitella*; XP_021190483.1, *Helicoverpa armigera*; XP_021700879.1, *Aedes aegypti*; XP_023939227.1, *Bicyclus anynana*; XP_032519693.1, *Danaus plexippus*.

### Immuno-fluorescent staining of sperm bundles

The anit-BmVasa antibody (BmVasa-R1) was described previously [53,66]. Immunofluorescence staining experiments using anit-BmVasa antibody were performed using spermatocytes and sperm bundles isolated from excised testes (fifth instar larvae stage to adult stage). The collected sperm were fixed for 1 h (Beyotime). After two times washing in PBS, samples were incubated in the primary antibody blocking solution mixture overnight at 4□ (PBS containing 0.1% Triton X-100 and 1% BSA (bovine serum albumin)). After two washes in PBS, samples were incubated with the secondary antibody, TRITC Phalloidin (YEASEN), and Hoechst (Beyotime) for 1 hr at room temperature. Samples were washed three times with PBS and subsequently mounted in the antifade medium (YEASEN). All images were taken on an Olympus BX53 microscope.

Antibodies and their concentrations were used as follows: BmVasa-R1, 1:200; Alexa Fluor^®^ 488 AffiniPure Goat Anti-Rabbit IgG (H+L), 1:500 (Thermofisher); TRITC Phalloidin, 1:500 (YEASEN).

### Western blotting and antibodies

Antibodies and their concentrations were used as follows: BmVasa-R1, 1:1000; monoclonal mouse anti-α-actin, 1:1000 (EpiZyme); HRP Goat Anti-Rabbit IgG (H+L), 1:1000 (EpiZyme); HRP Goat Anti-Mouse IgG (H+L), 1:1000 (EpiZyme). The Immobilon™ Western Chemiluminescent HRP substrate Kit (Millipore) was used to detect the protein signal.

### RNA-sequencing (RNA-seq) analysis

Total RNA from the testes and sperm bundles at day four in the fifth larval stage animals was extracted from six individual animals of WT, *ΔBmPrmt5*, and *ΔBmVasa*, and mixed together. The sperm bundles were released by tearing testis in PBS buffer. The mRNA was firstly enriched, and then fragmented and used for cDNA synthesis and library construction. The library was sequenced using Illumina 2000 platform. The raw data were qualified, filtered, mapped, and quantified (FastQC, Trimmomatic, Bowtie2, Rsem) to the reference silkworm genome database (http://silkbase.ab.a.u-tokyo.ac.jp/cgi-bin/index.cgi) [67–70], and mRNA abundance was normalized with Deseq2. The calculated gene expression levels were then used to compare gene expression differences among samples (Y/X). Differentially expressed genes (DEGs) were screened based on the Poisson Distribution Method with a false discovery rate (FDR) < 0.05 and the absolute value of log_2_(Y/X) > 1. Enrichment analyses of DEGs were conducted using the Kyoto Encyclopedia of Genes and Genomes (KEGG) database (org.bmor.eg.db). The visualization was processed by using R packages (ggplot2, pheatmap).

### Statistical analysis

All experiments in this study were performed with at least three replicates (except the RNA-seq). All data are expressed as the mean ± standard error (SEM). Differences between groups were examined using either a two-tailed Student’s t-test or a two-way analysis of variance. All statistical calculations and graphs were made with GraphPad Prism version 9.

## Supporting information

**S1 Table. Primers used in this work.**

**S1 Fig.**
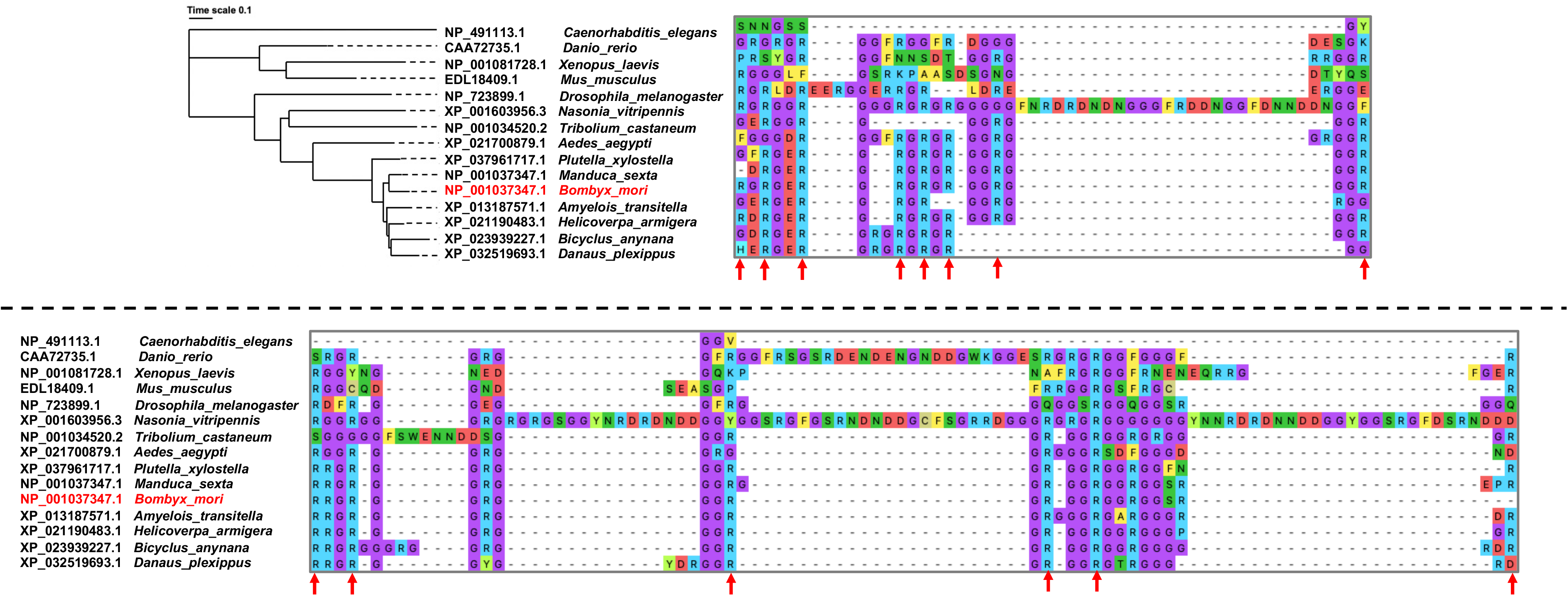
Phylogenetic and amino acid alignment analysis of Vasa protein in different species. The images of aligned sequences are predicted as symmetric dimethylation arginine residues (sDMA), which are denoted by red arrows.

**S2 Fig.**
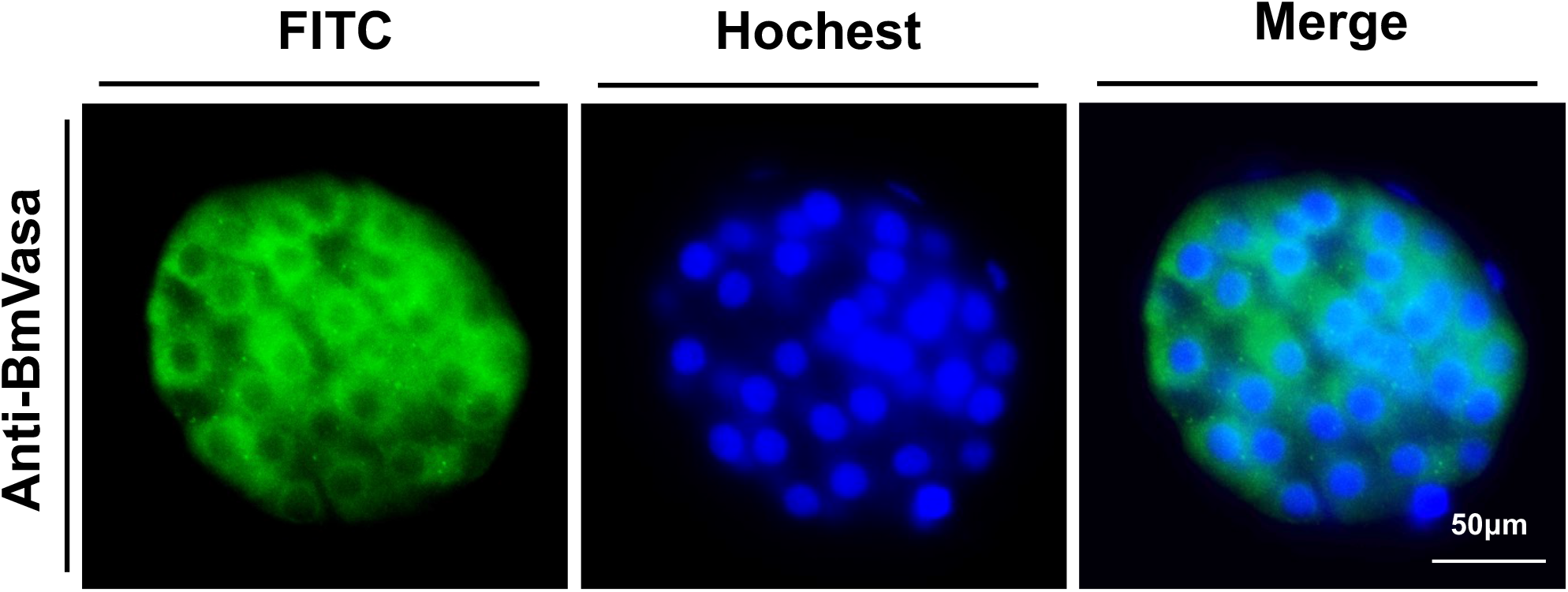
Representative immunofluorescence staining images of spermatocytes at day four of the fifth larval instar. Green, BmVasa; Blue, Hoechst. Scale bar, 50 μm.

**S3 Fig.**
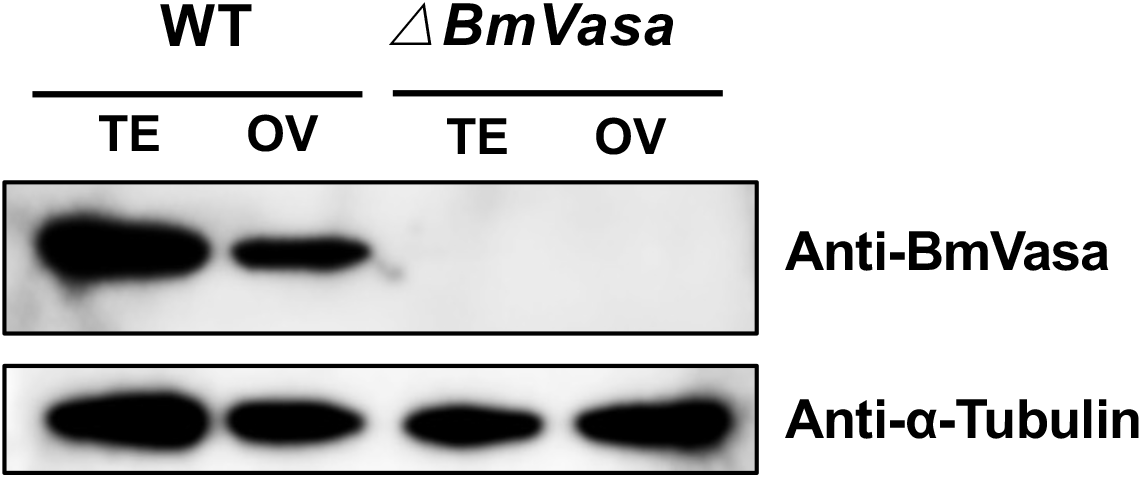
Western blot analysis of BmVasa protein in *ΔBmVasa*. TE and OV dentote testis and ovary, respectively.

**S4 Fig.**
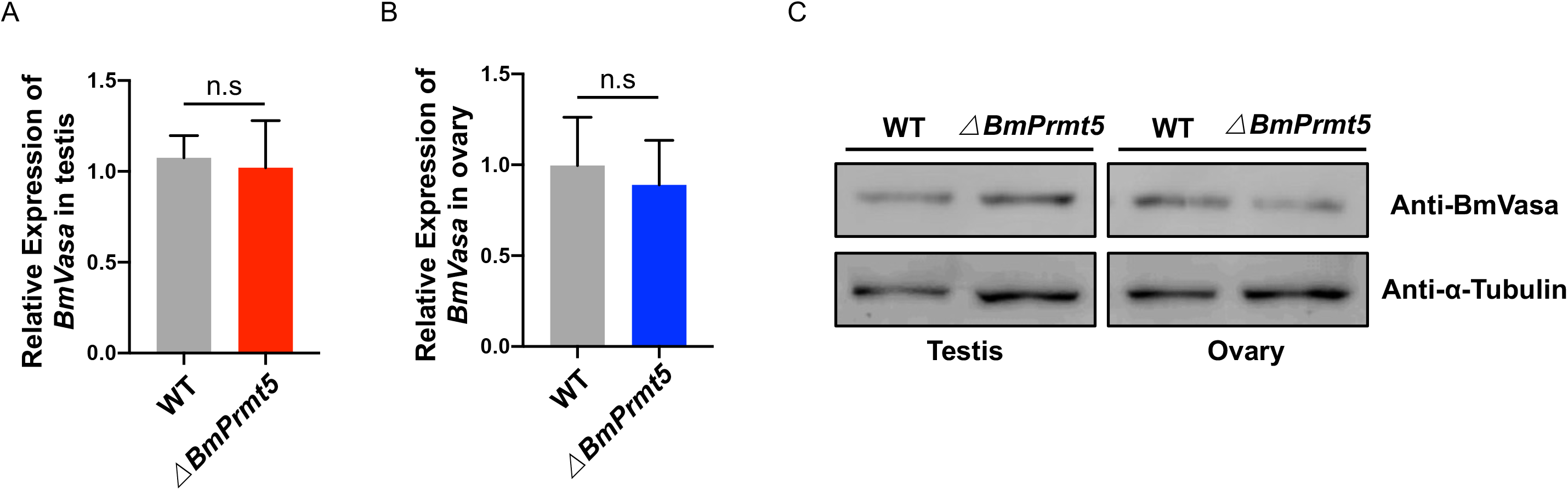
Detection of *BmVasa* transcript and BmVasa protein levels in *ΔBmPrmt5* mutant. (A and B) qRT-PCR analysis of *BmVasa* transcript in *ΔBmPrmt5* testis and ovary; (C) Western blot analysis of BmVasa protein levels in *ΔBmPrmt5* testis and ovary.

**S5 Fig.**
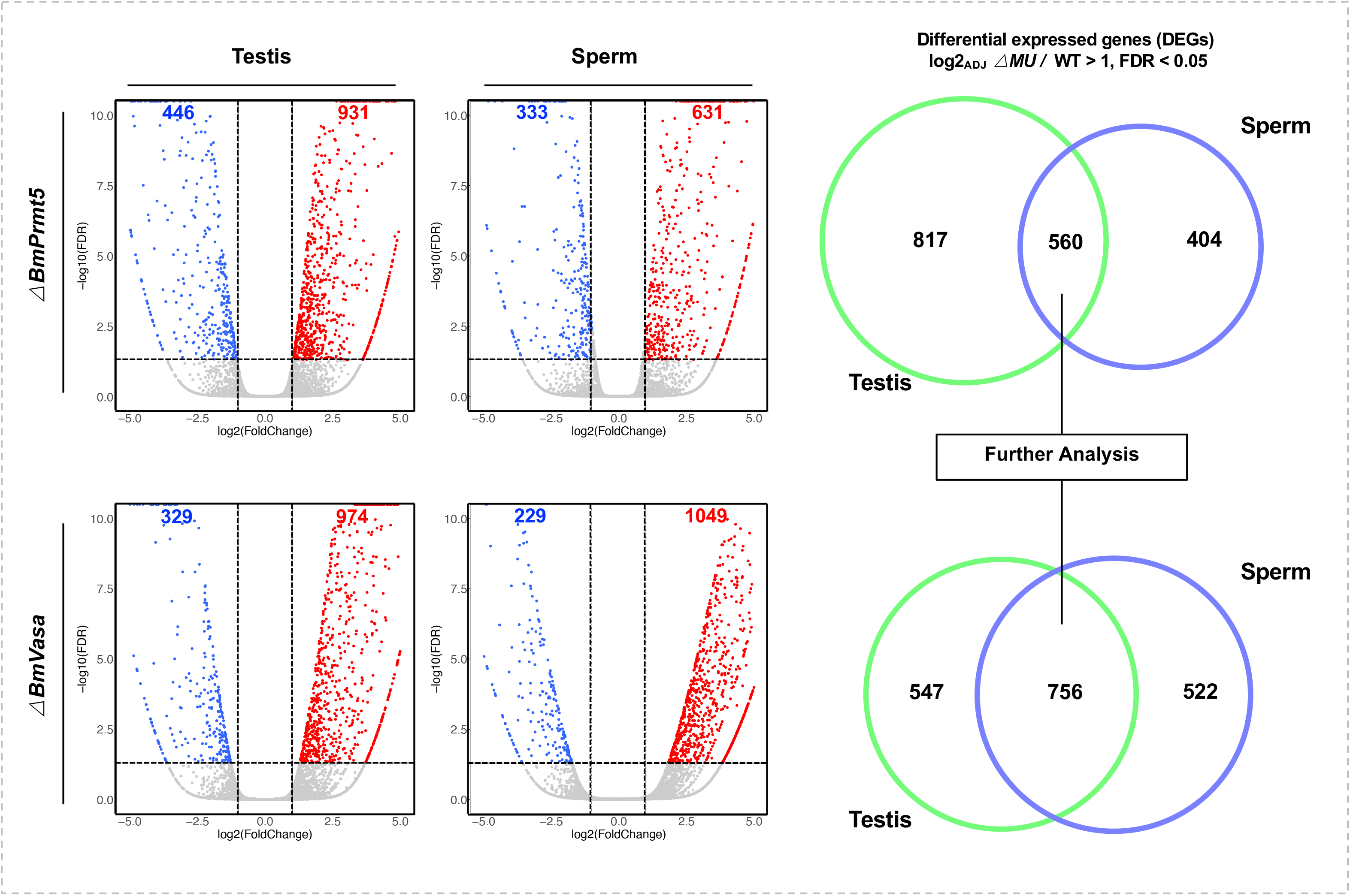
A global overview of the gene expression levels among all the libraries and volcano plots for DEGs of *ΔBmPrmt5* and *ΔBmVasa* samples. Volcano plots of differential expressed genes (DEGs) in *ΔBmPrmt5* and *ΔBmVasa* testis and sperm samples (Left panel). Red and blue represent DEGs as a fold change > 1 and FDR < 0.05. Venn diagram analysis (right panel) shows the DEGs commonly and specifically regulated in testis (green) and sperm (blue) samples of *ΔBmPrmt5* and *ΔBmVasa*. The sperm-specific DEGs in *ΔBmPrmt5* and *ΔBmVasa* were used for further analysis.

## Acknowledgments

The authors thank Prof. Hong-Quan Yang for proofreading the manuscript. We also thank Lianyan Jing and Shanshan Wang (Core Facility Centre of the Institute of Plant Physiology and Ecology) for their technique support. This research was supported by the National Science Foundation of China (32021001) and the Strategic Priority Research Program of the Chinese Academy of Sciences (Grant No. XDPB16).

## Author Contributions

**Conceptualization:** Xu Yang

**Data curation:** Xu Yang, Dongbin Chen, Shirui Zheng, Meiyan Yi, Yongjian Liu

**Formal analysis:** Xu Yang

**Funding acquisition:** Yongping Huang

**Investigation:** Xu Yang

**Methodology:** Xu Yang

**Resources:** Yongping Huang

**Supervision:** Yongping Huang

**Visualization:** Xu Yang

**Writing – original draft:** Xu Yang

